# Biodiversity of Magnetotactic Bacteria in the Freshwater Lake Beloe Bordukovskoe, Russia

**DOI:** 10.1101/682252

**Authors:** V. V. Koziaeva, L. M. Alekseeva, M. M. Uzun, P. Leão, M. V. Sukhacheva, E. O. Patutina, T. V. Kolganova, D. S. Grouzdev

## Abstract

According to 16S rRNA gene- or genome-based phylogeny, magnetotactic bacteria (MTB) belong to diverse taxonomic groups. Here we analyzed the diversity of MTB in a sample taken from the freshwater lake Beloe Bordukovskoe near Moscow, Russia, by using molecular identification based on sequencing of the 16S rRNA gene and *mamK*, a specific marker gene for these bacteria. A protein encoded by the *mamK* gene is involved in magnetosome chain arrangement inside the cell. As a result, six operational taxonomic units (OTUs) of MTB were identified. Among them, OTUs affiliated with the phylum *Nitrospirae* were predominant. ‘*Ca*. Etaproteobacteria’ and *Alphaproteobacteria* represented the minor groups of MTB. We also identified a novel MTB belonging to the family *Syntrophaceae* of the *Deltaproteobacteria* class. Using a combination of fluorescence and transmission electron microscopy, the bacteria belonging to these new MTB groups were visualized. Electron microscopy revealed that *Syntrophaceae* MTB were rod-shaped and synthesized elongated magnetosomes, arranged as a disorganized cluster. Among the *Nitrospirae* group, two groups with vibrioid cell shape and one group of ovoid-shaped bacteria were identified, all of which had elongated magnetosome crystals consisting of magnetite.

---

Prokaryotes capable of directed active movement that is guided by geomagnetism are collectively called magnetotactic bacteria (MTB) (Blakemore, 1975). MTB are physiologically, morphologically and phylogenetically diverse, sharing only the ability to synthesize special organelles termed magnetosomes. Magnetosomes consist of nanometer-size magnetite (Fe_3_O_4_) or greigite (Fe_3_S_4_) crystals surrounded by a lipid bilayer membrane containing proteins specific to the organelle. Magnetosomes are often assembled into chains inside the cell. MTB evolved the ability to conduct a special type of movement called magnetotaxis, which is based on orientation relative to magnetic field lines (Faivre and Schuler, 2008). Specific genes involved in magnetosome formation were found in the genomes of all MTB. There are currently nine highly conserved primary genes in magnetite- and greigite-producing MTB (*mamA, mamB, mamM, mamQ, mamO, mamI, mamP, mamK*, and *mamE*) (Lefèvre and Bazylinski, 2013). One of them, the *mamK* gene, encodes the actin-like protein MamK, which is responsible for the ordered arrangement of the magnetosome chain in the cell (Uebe and Schüler, 2016).

MTB are found within the phyla *Proteobacteria, Nitrospirae, Planctomycetes*, the candidate phyla *‘Omnitrophica’* and *‘Latescibacteria’* (Dziuba et al., 2016; Lefèvre and Bazylinski, 2013; Lin et al., 2017a; Lin and Pan, 2015). ‘*Ca*. Etaproteobacteria’ and *Alphaproteobacteria* are the most commonly observed and extensively characterized types of MTB (Lefèvre and Bazylinski, 2013; Monteil et al., 2018). Many of cultured marine and freshwater MTB belong to the class *Alphaproteobacteria*, including various vibrios and spirilla affiliated to the genera *Magnetospirillum, Magnetovibrio, Magnetospira*, and *Terasakiella* (Lefèvre and Bazylinski, 2013; Monteil et al., 2018). Numerous magnetotactic cocci and rods are affiliated to the order *Magnetococcales* of the class ‘*Ca*. Etaproteobacteria’ (Koziaeva et al., 2019). The two *Deltaproteobacteria* orders, *Desulfovibrionales* and the *Desulfobacteriales*, also contain magnetotactic members. Cultured strains *Desulfovibrio magneticus* RS-1, FH-1, ML-1, AV-1, ZZ-1, and an uncultured bacterium WYHR-1 belong to the order *Desulfovibrionales* (Lefèvre and Bazylinski, 2013; Li et al., 2019). The order *Desulfobacteriales* contains the strains *Desulfamplus magnetovallimortis* BW-1^T^ and SS-2, as well as several uncultured multicellular magnetotactic prokaryotes (MMP) (Abreu et al., 2007; Leão et al., 2017; Lefèvre and Bazylinski, 2013).

Axenic cultures of MTB of the *Nitrospirae* phylum have not yet been obtained. According to Parks et al., the most well-known members of this group form the candidate family ‘Magnetobacteraceae’. The family *‘Ca*. Magnetobacteriaceae*’* currently includes several candidate genera, which accommodate only MTB: *‘Ca*. Magnetobacterium, ‘*Ca*. Magnetoovum*’*, and *‘Ca*. Magnetominusculus*’* (Lefèvre and Bazylinski, 2013; Lin et al., 2011, 2017b). Most of MTB belonging to the *Nitrospirae* phylum were found in mesophilic freshwater habitats, with the exception of two representatives, WGC and BGC, the recently described giant cocci with multiple chains of magnetosomes (Qian et al., 2019), and multiple sequences retrieved from the Yelow Sea in China (Xu et al., 2018). A thermophilic MTB named *‘Ca*. Thermomagnetovibrio paiutensis*’* HSMV-1 was also described. It is closely related to cultured *Thermodesulfovibrio* spp.

Despite the high diversity of MTB found in environmental samples, they are difficult to isolate in axenic culture. Therefore, culture-independent techniques are indispensable in research on these bacteria. Members of the phyla *Proteobacteria* and *Nitrospirae* have been studied extensively. However, analyses of metagenomic datasets revealed the presence of MTB within the phyla *Planctomycetes* and ‘*Ca*. Latescibacteria’ (Lin and Pan, 2015; Lin et al., 2017a). This finding indicates that the biodiversity of MTB is much broader than is currently known. Therefore, modifying and developing the strategies for investigation offers great promise towards identifying MTB groups. Another challenge in MTB biodiversity investigation is detailed characterization of cell morphology and crystallographic properties of their magnetosomes. A recently developed method of coordinated fluorescence *in situ* hybridization and electron microscopy (FISH-TEM) successfully linked the phylogeny with cell ultrastructure and magnetosome morphology of several uncultured MTB strains, including the Gammaproteobacteria strain SHHR-1, Deltaproteobacteria strain WYHR-1 and ‘*Ca*. Magnetaquicoccus inordinatus’ UR-1 (Li et al., 2017, 2019; Koziaeva et al., 2019).

Here we describe a diversity of MTB in the lake Beloe Bordukovskoe based on a novel isolation approach that does not depend on cell motility. For phylogenetic identification of collected cells, we used a universal primer system on the marker gene *mamK* and analysis of the 16S rRNA gene sequences. To characterize the morphology of cells and magnetosomes of bacteria of retrieved OTUs, the FISH-TEM method was applied.

## MATERIALS AND METHODS

### Sampling and MTB enrichment

Water and sediment samples were collected at the freshwater lake Beloye Bordukovskoe, Shatura District, Russia (55°37’56”N, 39°44’38”E). Samples with a 1 : 2 sediment : water ratio were used to create a microcosm (3 L) and incubated in the dark at room temperature for one year. The pH value and salinity were measured using an Acvilon pH meter (Russia) and a handheld portable refractometer, respectively. The elemental composition of organic carbon (C%) in the sediment was measured using a Flash 1112 elemental analyzer coupled with a Thermo-Finnigan Delta V Plus isotope mass spectrometer (Thermo Fisher Scientific, United States) at the Core Facility of the Severtsov Institute of Ecology and Evolution, Russian Academy of Sciences.

Magnetic collection of MTB was performed using the MTB-CoSe (magnetotactic bacteria column separation) method developed in our laboratory (Fig. 1). Initially, sediment cores (15 mm in diameter) were taken from three different sites of the microcosm and mixed. To desorb the cells from large clay particles, PBS buffer (final concentration 1.25×) and glass microbeads (2 g; 150– 200 μm diameter; Sigma-Aldrich, United States) were added to a 50-mL tube containing 20 mL of the sample (sediment + water in approximately 1 : 4 ratio). The mixture was then treated on a ThermoShakerTS-100 (Biosan, Latvia) for 15 min at 100 rpm. The homogenate was transferred to a Bunsen flask and filtered under vacuum through a paper filter to remove large soil particles. The filtrate was transferred to 50-mL tubes and centrifuged (Eppendorf Centrifuge 5804R) for 10–15 min at 8000 rpm. Most of the supernatant was discarded, and 2 mL was retained for cell resuspension. Next, a miniMACS column (Miltenyi Biotec, Germany) was washed with 1.25× PBS buffer and placed on a magnet, and the cell mixture was applied to the column. MTB cells that adsorbed onto the column were washed 4–5 times with 2 mL 1.25× PBS buffer until no cells were present in the washing liquid (controlled by light microscopy). Next, the column was removed from the magnet, and the MTB cells were eluted twice with 100 μL 1.25× PBS buffer into a clean 1.5-mL tube. For samples with high cell concentrations, the eluate was reapplied onto the column, which was rinsed to completely remove nonmagnetic cells. The presence of MTB cells and nonmagnetic bacteria was assessed by microscopy of 5-μL aliquots of eluted liquid. The MTB sample was divided into two parts. One part was used to isolate DNA for metagenome sequencing and creating libraries of the *mamK* and 16S rRNA gene sequences. The second part was fixed for FISH/TEM studies. Genomic DNA was extracted using the DNeasy PowerSoil kit (Qiagen, Netherlands) according to the manufacturer’s instructions.

**Figure 1.**
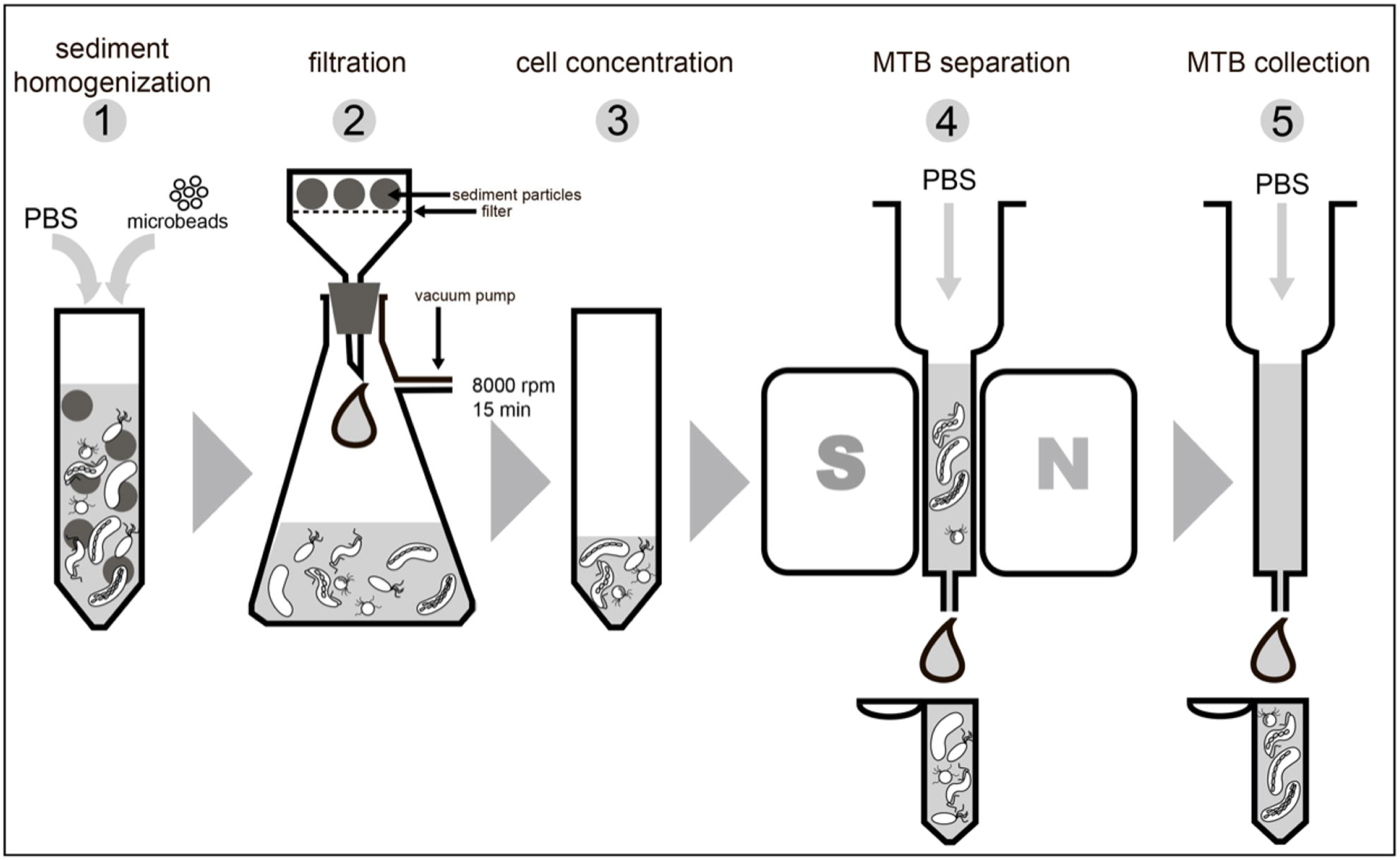
Scheme of MTB-CoSe separation procedure that allows collection of magnetotactic components directly from the sediment samples.

### Amplification, cloning and sequencing fragments of genes encoding 16S rRNA and *mamK*

A fragment of the *mamK* gene was obtained using nested PCR with two pairs of degenerate primers: external (mamK_79F and mamK_577R) and internal (mamK_86F and mamK_521R) (Table 1). The reaction buffer (25 μL) had the following composition: 5 μL Mas^CFE^TaqMIX-2025 buffer (Dialat Ltd., Russia), 0.5 pmol/μL each primer, 0.04% BSA, and 10–50 ng of a DNA template. The temperature-time profile of the reaction was as follows: initiation, 3 min at 95 °C; 4 cycles of 30 s at 95 °C, 40 s at 58 °C, and 1 min at 72 °C; 36 cycles of 30 s at 95 °C, 40 s at 52 °C, 1 min at 72 °C; and a final cycle of 7 min at 72 °C. Amplification of the 16S rRNA gene sequences was performed using the universal primers 27F and 1492R (Lane, 1991). Clonal libraries of the *mamK* and 16S rRNA gene fragments were obtained and sequenced as previously described (Kozyaeva et al., 2017). The target insert of the *mamK* gene was sequenced using the primer M13F, and 16S rRNA gene was sequenced using the primers 341F, 530F, 1114F and 519R (Sambrook et al., 1989).

**Table 1.**
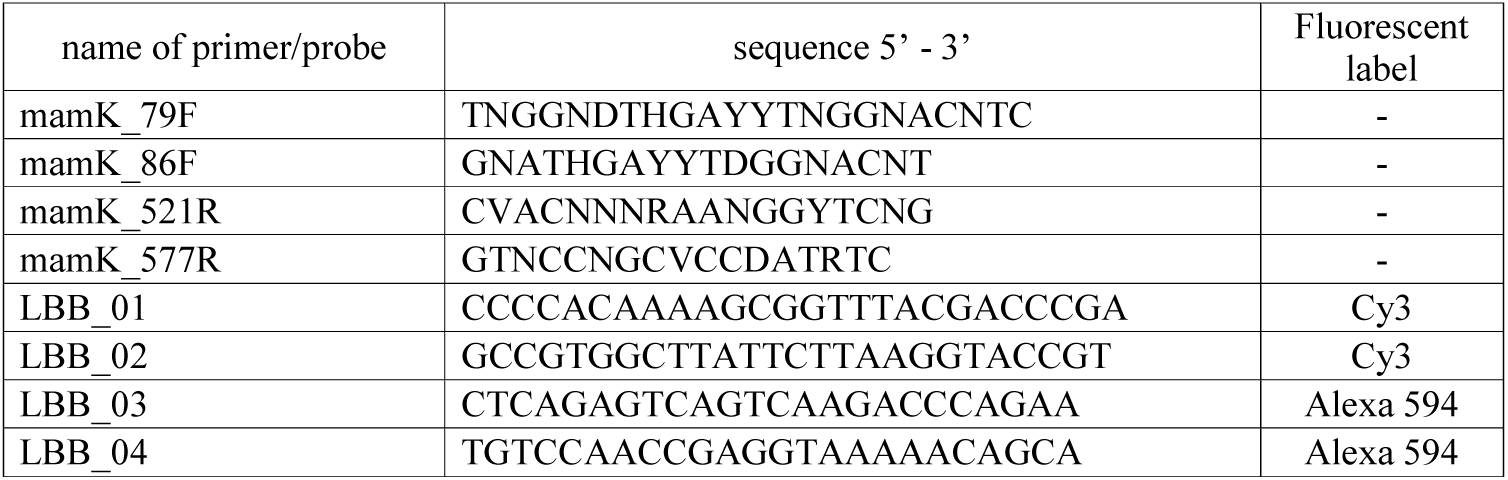
The list of primers and probes designed for *mamK* gene amplification and for FISH-TEM analysis, respectively.

### Phylogenetic analysis

Nucleotide and amino acid sequences were aligned using MAFFT (Katoh and Standley, 2013). Obtained MamK and 16S rRNA gene sequences were grouped in OTUs using the identity threshold of 97%. Phylogenetic analysis was performed using the IQ-TREE program (Nguyen et al., 2015) with selection of an evolutionary model using ModelFinder (Kalyaanamoorthy et al., 2017) and estimation of branch supports using UFBoot2 (Hoang et al., 2017). The MamK and 16S rRNA gene sequences representing OTUs identified in this study were deposited in GenBank under the accession numbers MK636828-33 and MK63285-90, respectively.

### Light and electron microscopy

Morphology of the cells collected after magnetic separation using MTB-CoSe was examined using an Eclipse E200 light microscope (Nikon, Japan). For conventional transmission electron microscopy (TEM), magnetically enriched cell suspensions were deposited on Formvar-coated 300-mesh copper grids, washed with distilled water and imaged using a Morgagni TEM (FEI, United States) operated at 80 kV.

### Fluorescence in situ hybridization/transmission electron microscopy (FISH-TEM)

For fluorescence in situ hybridization (FISH), collected MTB cells were fixed in 3% paraformaldehyde for 1.5 h. For the association of the retrieved 16S rRNA gene sequences with different MTB morphotypes, a drop of sample was added to Formvar-coated center-marked or index copper grids. A thin layer of carbon was sputtered onto the grids (Balzers CED-030/Baltec) atop each sample to provide stabilization. FISH was performed on each grid using the conditions and buffers described by Pernthaler et al. (2001) and 30% formamide in the hybridization buffer. After this procedure, the grids were stained with 0.1 μg/mL 4,6-diamidino-2-phenylindole (DAPI) for 10 min, placed between a glass slide and cover glass, and observed with a Zeiss AxioImager microscope equipped with an AxioCam Mrc (Zeiss, Germany).

Probes used for FISH were designed based on the 16S rRNA gene sequences of retrieved OTUs (Table 1). In addition, a mix of EUB388I, EUB388II and EUB388III bacterial universal probes labelled with Alexa 488 was used as a control (Daims et al., 1999).

After performing FISH, the same grids were examined by TEM, and images were taken in the same region where hybridization with the specific probe occurred.

### Metagenome binning and analysis

The metagenome 3300021602 obtained from the IMG database was binned using three different tools, MaxBin2 (Wu et al., 2014), MyCC (Lin and Liao, 2016), and Busy Bee Web (Laczny et al., 2017) prior to dereplication and refinement with the DAS Tool (Sieber et al., 2018), which performs a consensus binning to produce the final bin set. Completeness and contamination rates were assessed using CheckM v. 1.0.12 (Parks et al., 2015) with the ‘lineage wf’ command and default settings. Magnetosome island genes were found using local BLAST compared with reference sequences of magnetotactic bacteria.

## RESULTS

### Collection of MTB from a sediment sample

Lake Beloye Bordukovskoe is a round lake with a maximum depth of 21 m and is possibly of glacial origin. The lake has transparent water with a light brown color and visibility of up to 8 m. The brown color is due to water flow from the nearby peat bog. The pH of the microcosm water was 6.3 and salinity was 1% during sampling. We used an MTB-CoSe method to enrich MTB from the microcosm without inducing magnetotaxis. Immediately after magnetic separation, light microscopy examination revealed the presence of 1.12 ± 0.06 × 10^6^ MTB in 20 mL of the total sample. Vibrios were the most common MTB morphotype, and ovoid cells, which accounted for about 30% of the magnetic fraction, were also detected. Nonmagnetic cells were also present in small quantities (<1%).

### Phylogenetic analysis of the MamK sequenses

After the removal of chimeras and low-quality reads, the clone library of the *mamK* gene fragments consisted of 346 sequences. Based on the derived MamK sequences, phylogenetic analysis was conducted that yielded six OTUs (Table 2; Fig. 2a). MamK_LBB_01, MamK_LBB_02, and MamK_LBB_03 were the most well-represented OTUs. On the phylogenetic tree, these OTUs clustered together with the MamK sequences of strains of the *Nitrospirae* phylum. The dominant OTU MamK_LBB_01 was represented by 87% of all clones and formed a separate clade within the phylum *Nitrospirae*. The minor groups were OTUs MamK_LBB_05 and MamK_LBB_06, which clustered with the MamK sequences of MTB belonging to the *Magnetococcales* and *Rhodospirillales* orders.

**Table 2.**
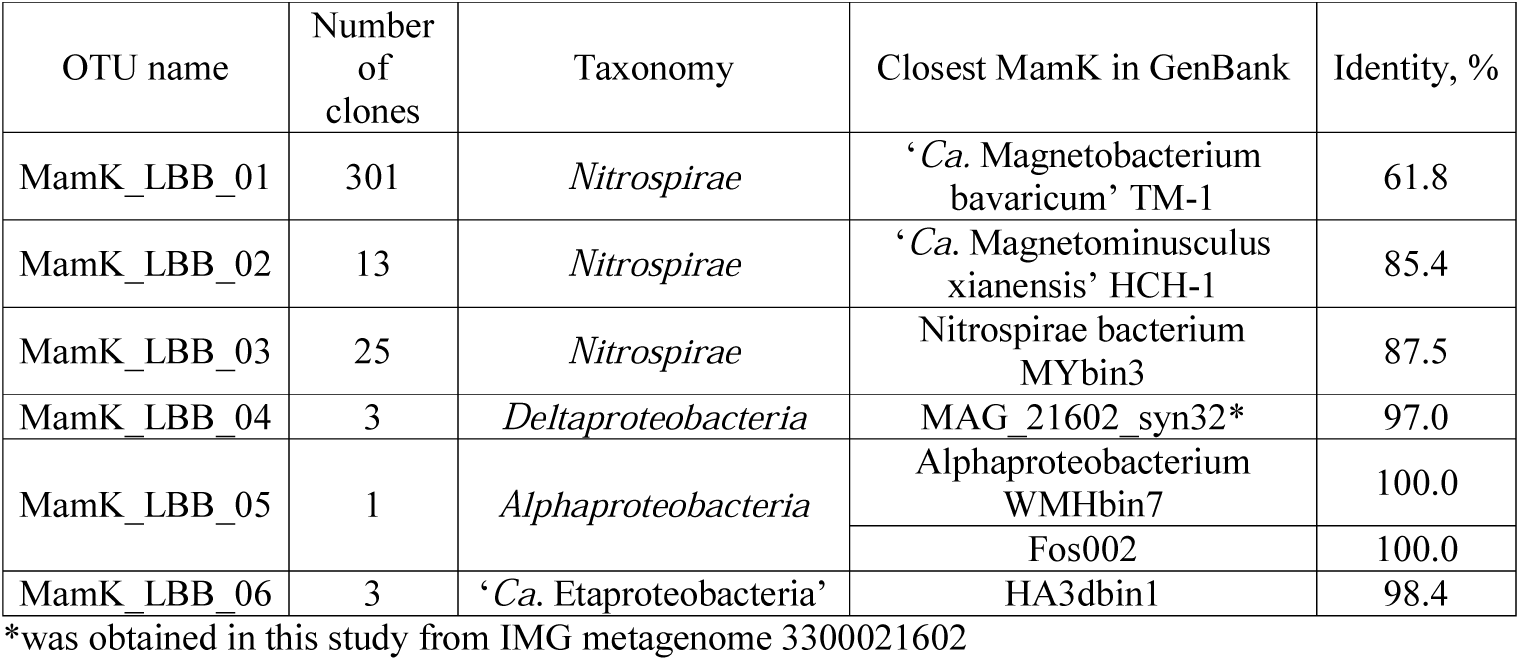
Operational taxonomic units (OTUs) of MamK sequences retrived in this study and closest representatives of MTB

**Figure 2.**
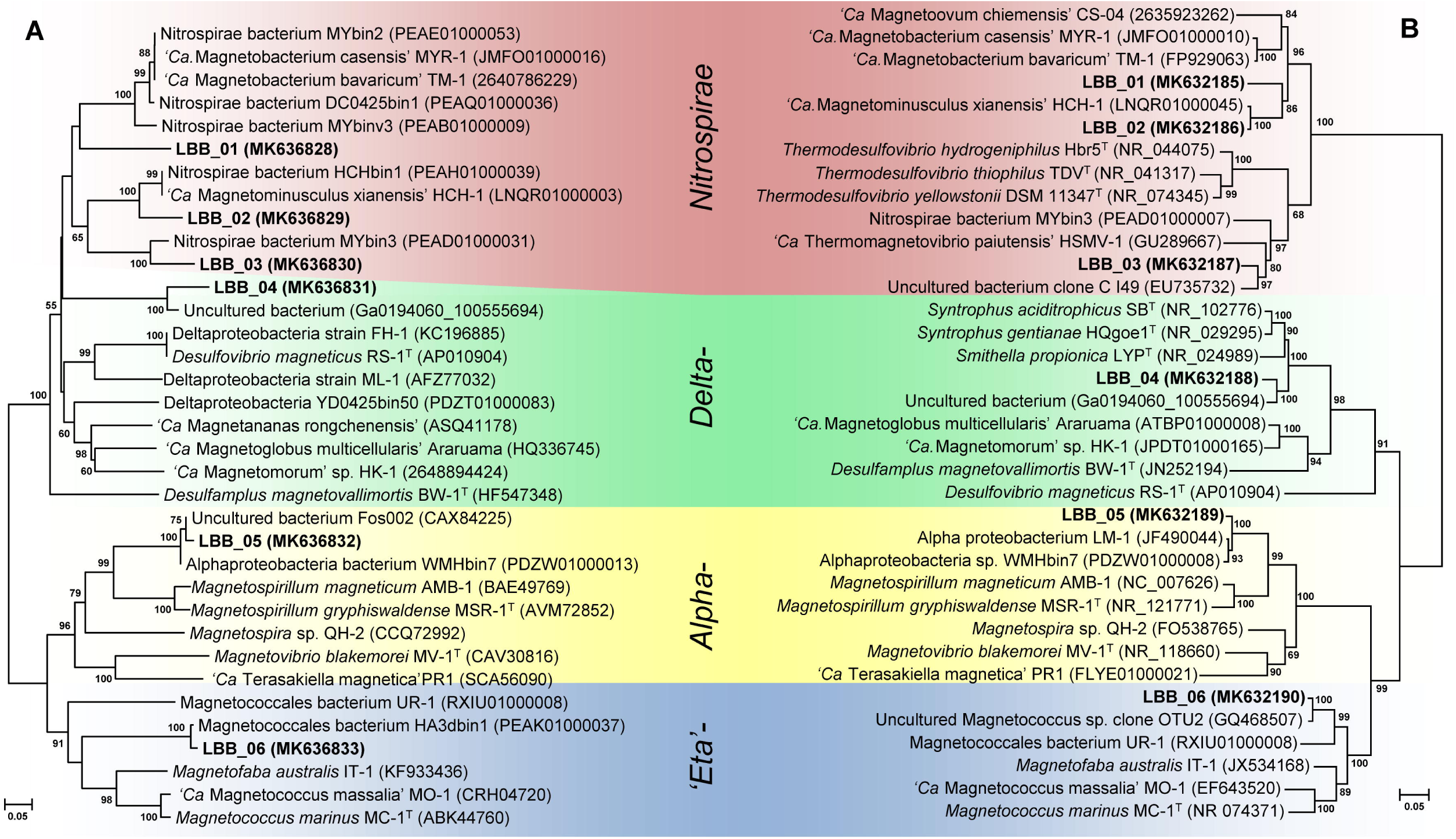
Maximum-likelihood phylogenetic trees based on magnetosome associated protein MamK (285 amino acid sites) reconstructed with evolutionary model LG+I+G4 (a); and 16S rRNA gene sequences (1,399 nucleotide sites) reconstructed using the evolutionary model GTR+F+I+G4 (b).

One interesting finding was the presence of the OTU MamK_LBB_04, which was clearly divergent from all other currently known MamK sequences. The level of similarity to the closest MamK of the *Nitrospirae* bacterium MYbin3 was 69.2%, with 45% coverage. We could not attribute this OTU to any of the known taxonomic groups containing MTB due to the lack of related reference sequences. Therefore, we decided to conduct a search for possible homologues in the metagenomic dataset available in the public databases IMG and NCBI. After searching for the nearest homologues for the OTU MamK_LBB_04, a protein with a high sequence similarity (97.0%) was found in the IMG database. This MamK was recovered from the metagenome 3300021602, which was obtained from a boreal shield lake in Ontario (IISD Experimental Lakes Area). After binning of the metagenome, a bin (MAG_21602_syn32) containing the target *mamK* sequence was obtained. In addition to *mamK*, other magnetosome genes (*mamI, mamE, mamQ-1*, and *man2*) were found. The *man2* gene is located in the contig next to *mamK*. The OTU MamK_LBB_04 together with MamK, which is derived from MAG_21602_syn32, formed a phylogenetically distinct lineage between the *Nitrospirae* and *Deltaproteobacteria* branches. Taxonomy analysis of the bin MAG_21602_syn32 sequence from Genome Taxonomy Database showed that it belonged to the order ‘*Syntrophales’* and, according to NCBI, to the order *Syntrophobacterales*. Therefore the identified MamK could be associated with *Syntrophobacterales* species. MTB of this taxonomic group have not been previously detected. The 16S rRNA gene sequence of the received bin was also identified (acc. no. Ga0194060_100555694).

### Phylogenetic analysis of 16S rRNA gene sequenses

The assembled 16S rRNA gene clone library contained 282 sequences, which were classified into 6 different OTUs (Table 3). A phylogenetic tree was constructed and its topology was compared with the tree based on the MamK sequences (Fig 2b).

**Table 3.**
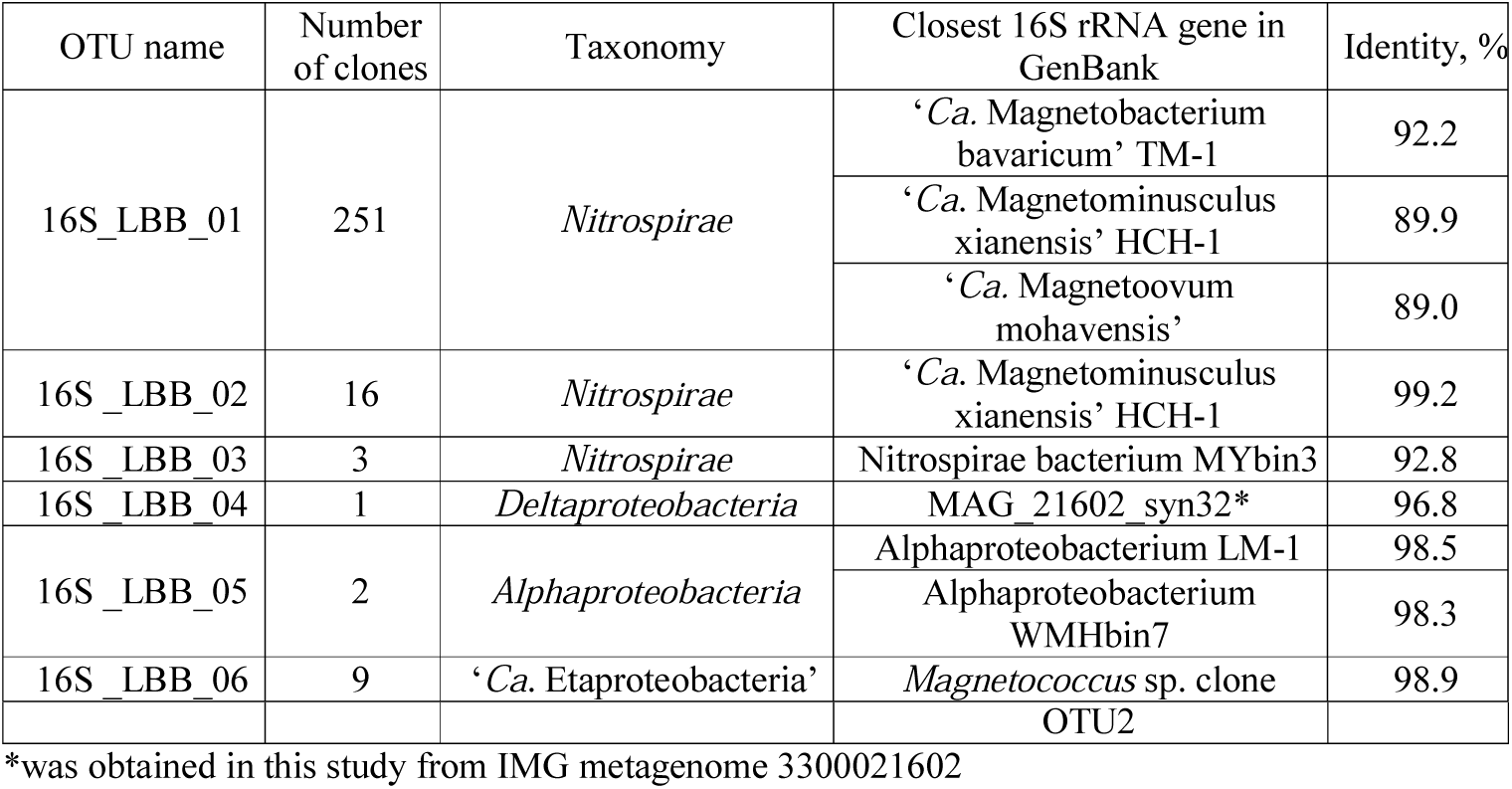
Operational taxonomic units (OTUs) of 16S rRNA gene sequences and closest representatives of MTB

Over 95% of the retrieved 16S rRNA gene sequences (OTU 16S_LBB_01□OTU 16S_LBB_03) clustered within the *Nitrospirae* phylum. A dominant group OTU 16S_LBB_01 formed a separate branch within the family ‘*Ca*. Magnetobacteraceae’ as well as on the phylogenetic tree based on the MamK sequences. OTUs 16S_LBB_06 and MamK_LBB_05 formed branches with bacteria of the *Magnetococcales* and *Rhodospirillales* orders, respectively, and represented 3.9% of the 16S rRNA gene clonal library.

OTU 16S_LBB_04 had 96.8% similarity with the 16S rRNA gene identified in the bin MAG_21602_syn32. On the 16S rRNA phylogenetic tree both sequences formed a separate branch with representatives of the family *Syntrophaceae* (order *Syntrophobacterales*). The level of similarity with the closest validly described species, *Syntrophus aciditrophicus* and *Smitella propionica*, was 93.5% and 91.2%, respectively, which indicates that OTU 16S_LBB_04 belongs to a novel genus within the family *Syntrophaceae*.

Topology of the trees was generally congruent, and both trees contained similar phylogenetic groups. It is possible that both the *mamK* and 16S rRNA gene sequences correspond to the same organism, but this hypothesis requires further verification. Due to the high level of similarity in 16S rRNA gene and MamK sequences, it is possible that OTU 16S_LBB_05, bacterium LM-1, and WMHbin7 are the same or closely related species of the *Rhodospirillales* order.

### Linking phylogeny to cell morphology by FISH-TEM

To link phylogeny with the MTB morphotype, FISH-TEM was carried out. Based on the target 16S rRNA gene sequences, the probes were designed to identify the morphology of the matching MTB by FISH-TEM (Table 1).

The LBB_01 probe specific for OTU 16S_LBB_01 hybridized only with vibrioid-shaped bacteria (Figs. 3a□3d). This group of MTB posessed a thick chain of anisotropic magnetosomes organized along the long axis of the bacterial cell. Our morphology identification analysis showed that bacteria of OTU 16S_LBB_02 were small ovoids and synthesized two bundles of bullet-shaped magnetosomes (Figs. 3e□3h). The observed cell shape and magnetosome organization was similar to that of *Nitrospirae* MTB fosmid MY3-11A (Lin et al., 2011). The LBB_03 probe, which was specific for OTU 16S_LBB_03, hybridized to a vibrioid cell containing anisotropic magnetosomes in a single chain (Figs. 3i□3l).

**Figure 3.**
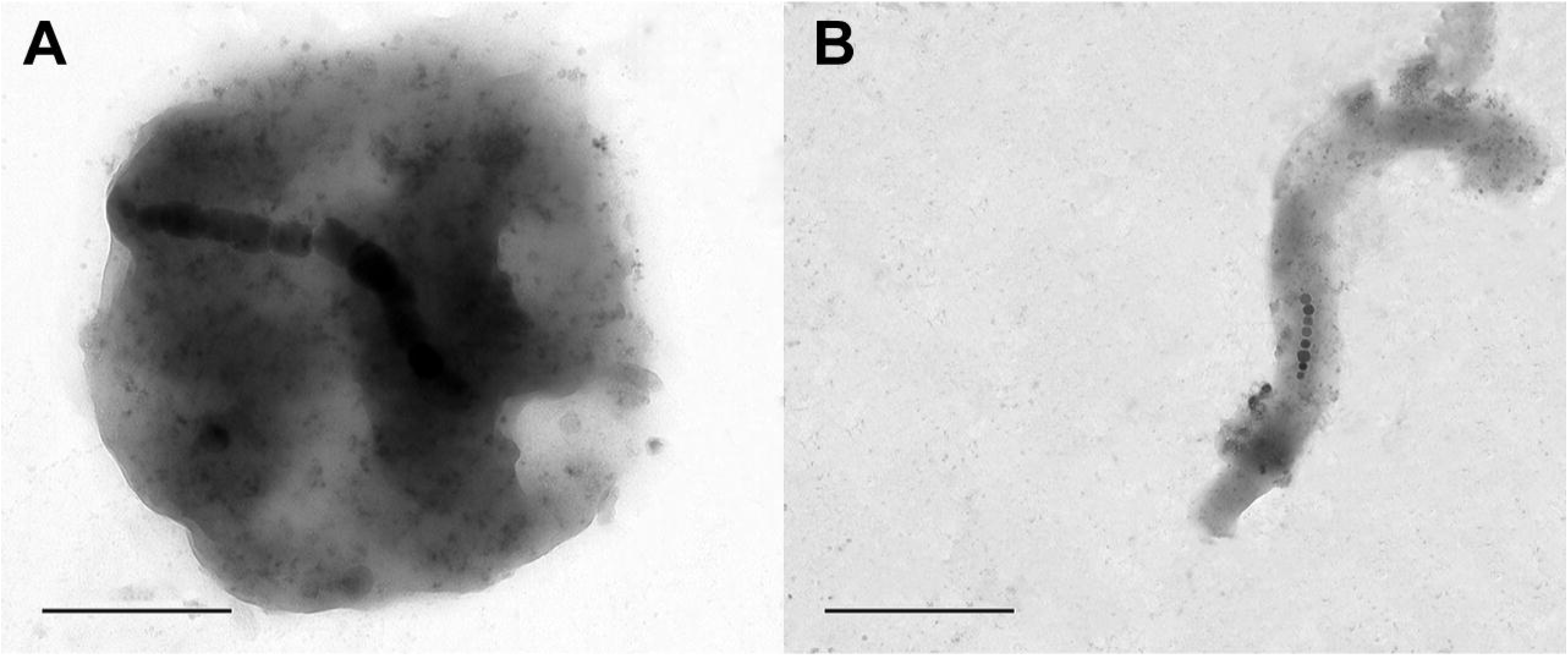
FISH-TEM images of MTB from environmental samples. The circles and arrows indicate the cells that did and did not, respectively, hybridize with the specific probe tested. Phase contrast image of magnetically enriched environmental samples on Formvar-coated TEM grids (a, e, i, and m); Bacteria present in the same region as imaged in a, e, i, and m, respectively, stained with DAPI (b, f, j, and n); Image of the same area captured in images b, f, j, and n after hybridization with the LBB_01, LBB_02, LBB_03, and LBB_04 probes, respectively (c, g, k, and o); TEM image of the same cell that showed hybridization (circles) with the specific probes in c, g, k, and o (d, h, l, and p).

To confirm that the identified OTU 16S_LBB_04 was related to magnetotactic bacteria, FISH-TEM analysis was conducted. The fluorescent probe specific to OTU 16S_LBB_04 identified a rod-shaped MTB with disorganized magnetosomes located close to the center of the bacterial cell (Figs. 3m□3p). Rod-shaped morphology is a common feature for bacteria of the *Syntrophaceae* family, which accommodates mesophilic bacteria inhabiting anoxic freshwater environments (Kuever et al., 2015).

For OTUs 16S_LBB_06 and MamK_LBB_05, FISH-TEM was not performed. However, a search of TEM images identified a coccoid cell with a single magnetosome chain (Fig. 4a). TEM images of spirilla with a chain of cubooctahedral magnetosomes were also observed (Fig. 4b).

**Figure 4.**
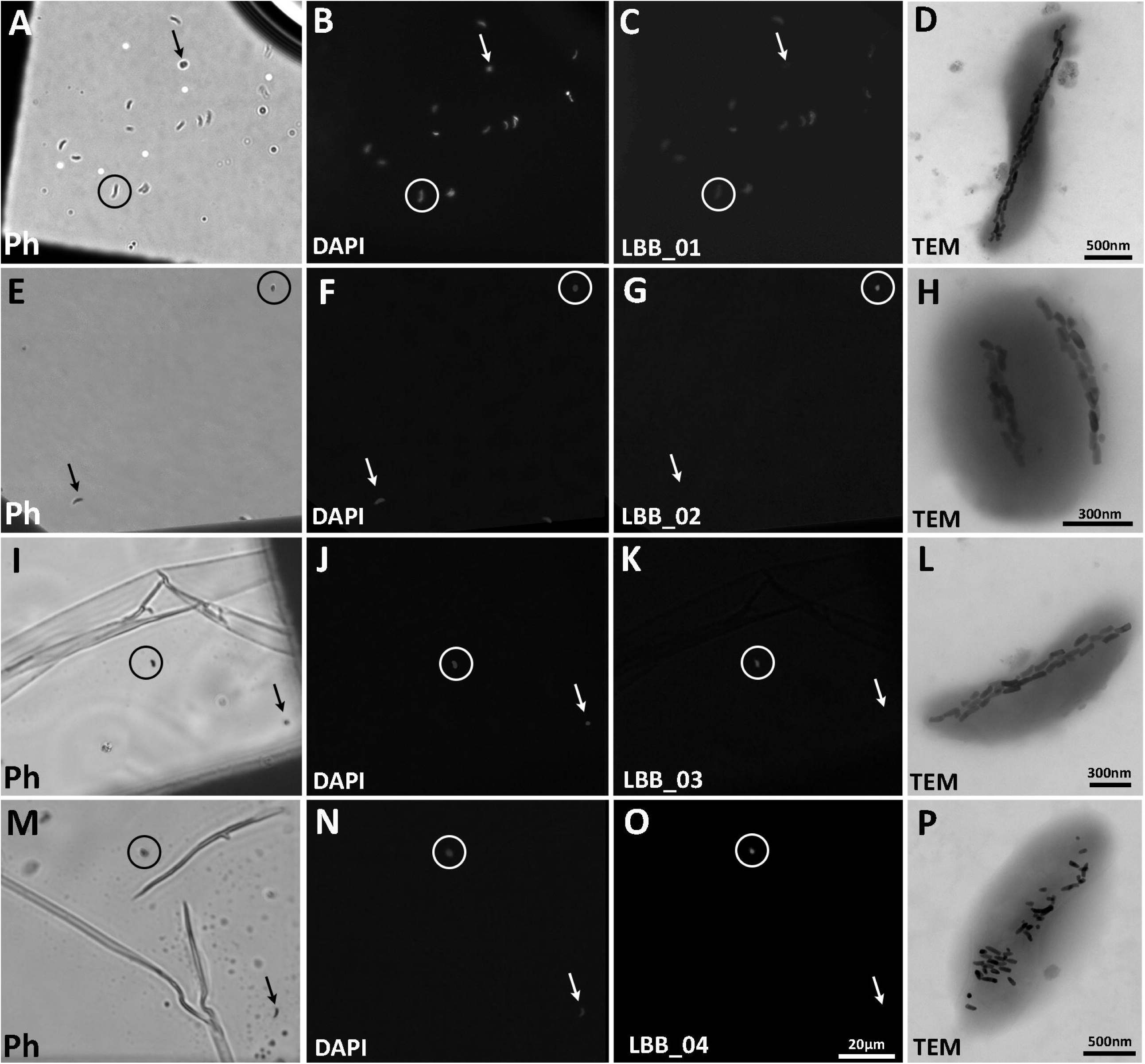
TEM images of magnetotactic coccus (a) and magnetotactic spirilla (b) identified in environmental samples. Bar 0.5 μm.

## DISCUSSION

Previous studies on the MTB diversity have mainly detected members of the *Alphaproteobacteria* and *‘Etaproteobacteria’* classes (Jogler et al., 2009; Lefèvre and Bazylinski, 2013; Kozyaeva et al., 2017; Lin et al., 2017a). However, we found that these two types of MTB were in fact minority groups in the Lake Beloye Bordukovskoe community. We found many representatives of the phylum *Nitrospirae*. Few studies have identified *Nitrospirae* as a dominant MTB phylum, including those at freshwater lakes Miyun (China) and Chiemsee (Germany) (Spring et al., 1993; Lin et al., 2011), as well as in the sediments from the continental shelf of the Yellow Sea (Xu et al., 2018). Differences between the studies may be due to a number of reasons. One main reason may be related to the procedure by which MTB are isolated from the environmental sample. *Alphaproteobacteria* and ‘*Ca*. Etaproteobacteria’ were usually the predominant classes or the only ones identified in the samples when a magnet was placed on the outside of the microcosm at the silt–water interface and then the cell suspension was purified using a “race-track” approach (Flies et al., 2005b; Lin et al., 2008; Postec et al., 2012; Dziuba et al., 2013; Pradel et al., 2016; Kozyaeva et al., 2017; Du et al., 2017;). Previous studies postulated the methods based on magnetotaxis to be highly dependent on the cell swimming ability of MTB (Lin et al., 2008; Lin and Pan, 2009; Xu et al., 2018). The intensity of the magnetic field, distance to the magnet, and aerotaxis also affect the enrichment efficiency. Within the oxic–anoxic interface (OAI), different MTB species show different vertical distributions. Magnetotactic cocci usually concentrate at the OAI (Flies et al., 2005a; Lefevre et al., 2011) and occur closer to the magnet during the first step of magnetic separation. Therefore, cocci of the ‘*Ca*. Etaproteobacteria’ class gain a significant advantage with this enrichment method. In the present study, we obtained sludge samples from a microcosm using a vertical tube, and as a result a larger amount of the anoxic zone was captured. Since the movement of MTB is directed not only by magnetic field, but also by aerotactic signals, it is possible that some MTB may not leave their anoxic zone during magnetic extraction of the cells from the microcosm sediment.

We cannot rule out the possibility that the dominance of members of the phylum *Nitrospirae* shown in this study was due to a limitation of the MTB-CoSe method. Representatives of the *Nitrospirae* phylum contain larger numbers of magnetic particles than other MTB, and since the procedure requires several washes, this may favor cells containing a large number of magnetosome crystals.

We also cannot exclude the possibility that during microcosm storage growth conditions were optimal for *Nitrospirae* MTB, rather than for magnetotactic cocci. The MTB community may change during incubation of the microcosm in the laboratory for a certain period of time (Flies et al., 2005b). It was previously shown that the numbers of magnetotactic cocci and spirilla significantly decreased over several months of the microcosm storage, whereas the numbers of the magnetotactic ovoid *Nitrospirae* bacterium LO-1 increased (Lefevre et al., 2011). Therefore, the distribution among the MTB phylogenetic groups may depend on the incubation time of the microcosm.

Magnetic separation methods represent robust tools for the investigation of MTB biodiversity and have been successfully used in numerous studies. However, any separation method can lead to a bias and thus may not reflect the real diversity of MTB in nature. As previously stated, the most acceptable approach for direct study of MTB biodiversity in natural samples is the one not involving magnetic separation (Lin and Pan, 2009). However, it is difficult to identify magnetotactic bacteria by direct amplification of their 16S rRNA gene sequences using universal primers. This is due to the fact that the ability to synthesize magnetosomes is not a taxonomic descriptor, and MTB and non-MTB may belong to the same taxonomic group (Lefèvre and Bazylinski, 2013). The absence of reference MTB strains in different taxonomic groups also makes it difficult to describe the diversity of MTB using the 16S rRNA gene sequence analysis. For example, the 16S rRNA gene sequences of magnetotactic ‘*Ca*. Latescibacteria’ and *Planctomycetes* have not been identified. Therefore, in this study we designed specific primers on the *mamK* gene sequences to overcome the problem of 16S rRNA gene profiling of the MTB community. We analyzed these nine genes of all known MTB from *Proteobacteria, Nitrospirae*, and ‘*Ca*. Omnitrophica’ phyla for primer system design. Based on our analysis, only *mamK* had a suitable length and conserved regions within all taxonomic groups for the design of a universal primer system. It was previously shown that some non-magnetotactic bacterial genomes contain homologues of MamK (Lefèvre et al., 2013). However, the *mamK* genes belonging to magnetotactic bacteria have the highest level of similarity with each other compared to the genes of non-magnetotactic bacteria. Therefore, phylogenetic analysis must be carried out extremely carefully to exclude potential non-magnetotactic MTB.

Over the past several years, the information on MTB has increased significantly, leading to the recent critical discovery of magnetotactic protists (Leão et al., 2019; Monteil et al., 2019). As the amount of genomic data increased, new taxonomic groups of MTB have been found, indicating that the biodiversity of magnetotactic bacteria remains underestimated. We expect that continued improvements in detection and separation techniques will lead to the identification of novel MTB in the future.

## ACKNOWLEDGMENTS

We thank Professor Josh D. Neufeld for permission to use metagenomic data (3300021602). We thank Dr. Svetlana Zhenilo for microscopy assistance and Dr. Vasil Gaisin for comments that improved an earlier version of this manuscript. We thank Jefferson Cypriano, CENABIO and Unimicro for helping with accsess to the TEM used in this work.

## COMPLIANCE WITH ETHICAL STANDARDS

The authors declare that they have no conflict of interest. This article does not contain any studies involving animals or human participants performed by any of the authors.

